# Optimized Blue Laser Priming Improves Lettuce Seed Germination and Early Seedling Establishment Under Reduced Water Availability

**DOI:** 10.64898/2026.08.14.744454

**Authors:** Ayesha Noor, Parviz Elahi

## Abstract

Continuous-wave (CW) 450 nm blue diode-laser irradiation was evaluated as a physical seed-priming treatment for improving lettuce (*Lactuca sativa* L.) germination and early seedling establishment under reduced water availability. Dry seeds were irradiated at 50–250 mW for 2 min and at 100 mW for 0.5–10 min; each condition included three independent Petri-dish replicates with 100 seeds per replicate, and the replicates were monitored for 72 h. Laser treatment produced a dose-dependent biological response. The optimized condition, 100 mW for 2 min, increased final germination from approximately 65–70% to 90–95%, increased the germination speed index, and promoted root elongation more strongly than shoot elongation. Longer exposures reduced germination and seedling growth. Under reduced water availability (0.5–6 ml per Petri dish), laser-treated seeds germinated earlier, maintained final germination of 83–93% compared with 57–72% in controls, and produced better-developed seedlings. The relative benefit increased as water availability decreased, indicating that optimized blue laser priming partially compensated for low water supply during germination and early establishment. These results identify CW blue laser priming as a contactless, chemical-free approach for improving lettuce seed performance and early seedling vigor under limited water availability.

## Introduction

Seed germination is one of the most decisive phases in the plant life cycle because the transition from a dormant seed to an actively growing seedling establishes the foundation for subsequent plant development, crop productivity, and agricultural sustainability.^1,2^ With the global population expected to approach nearly 8.5 billion, ^3^ improving the efficiency and uniformity of seed germination and early seedling development is increasingly important for supporting stable food production under changing environmental and resource-limited conditions. Conventional approaches used to enhance seed germination, including chemical priming treatments^4,5^ and mechanical or physical scarification, ^6,7^ have been widely applied. However, these methods can have limitations, including increased processing costs,^8^ potential environmental concerns, ^9^ and difficulty achieving uniform, precisely controlled treatment conditions. Therefore, non-chemical, energy-efficient, and controllable seed-stimulation techniques are attracting increasing research interest. Light-induced seed stimulation is one such physical priming strategy because light can regulate physiological and biochemical processes associated with germination and early seedling growth. Different optical sources, including sunlight, light-emitting diodes, continuous-wave (CW) lasers, and pulsed lasers, have been explored to improve germination behavior and plant establishment under controlled conditions.^10–16^ Among these approaches, laser-based seed stimulation is particularly attractive because lasers provide high directionality, monochromaticity, contactless treatment, and accurate control over irradiation parameters such as wavelength, output power, exposure duration, power density, fluence, and total delivered energy. These features make laser irradiation a highly tunable tool for seed priming, with the biological response adjustable by selecting an appropriate irradiation window. The wavelength-selective nature of laser irradiation is especially important in plant photobiology. ^17–19^ Seeds and developing seedlings contain photoreceptor systems that respond to specific spectral regions. For example, phytochromes, which exist in the Pr and Pfr forms, are mainly involved in red and far-red light responses, whereas cryptochromes and phototropins are primarily associated with blue-light perception.^11,20^ Therefore, a laser with a defined wavelength can provide targeted optical stimulation of selected light-responsive pathways. When applied at an optimized dose, laser irradiation may influence water uptake, enzymatic activation, metabolic regulation, cellular activity, protein synthesis, and the early development of root and shoot tissues. ^14,21^ However, because the laser-induced response depends strongly on wavelength, power, exposure time, and delivered energy, inappropriate irradiation conditions may produce neutral or inhibitory effects. Systematic optimization of laser parameters for each seed system is therefore essential for achieving beneficial germination and growth responses.

Low-level laser irradiation has been widely investigated as a physical seed-priming technique because it can induce measurable biological responses using relatively low energy input. Both CW and pulsed-laser systems have been reported to improve germination and early seedling performance in different plant species. For instance, Parveen et al. investigated the effects of different doses of low-power CW He-Ne laser irradiation on sunflower seeds and reported significant changes in germination-related enzymes, including amylase and proteases.^22^ Urva et al. studied *Moringa oleifera* seeds treated with low-power CW laser irradiation and found that laser exposure enhanced germination percentage, mean germination time, vigor index, root and shoot growth, and mineral accumulation, with the response depending on the applied laser energy.^23^ Podlesna et al. examined pre-sowing HeNe laser irradiation in pea seeds and showed that laser treatment improved germination rate and uniformity, increased amylolytic enzyme activity and indole-3-acetic acid content, and promoted plant height, leaf area, flowering, ripening, and yield-related traits. ^24^ Asghar et al. compared He-Ne laser treatment with magnetic-field pretreatment in soybean seeds and reported improvements in germination, seedling growth, biomass accumulation, and yield characteristics following laser exposure.^25^ Nadimi et al. evaluated red and dual-wavelength green/infrared laser biostimulation in flaxseeds and demonstrated that optimized laser treatment improved germination performance and seedling biomass, particularly in sub-optimally stored seeds with reduced germinability.^26^ Soliman and Harith investigated He-Ne laser irradiation in *Acacia farnesiana* seeds and reported that suitable irradiation conditions improved germination indices by affecting seed-coat permeability and modifying endogenous growth-regulating compounds such as GA, IAA, ABA, and phenols.^27^ Hasan et al. studied maize seeds exposed to red He-Ne and green Nd:YAG lasers and found that laser photobiomodulation enhanced field emergence, seedling growth, vigor index, and several growth-related traits.^28^ Sreckovi et al. examined the influence of diode and He-Ne lasers on corn and wheat seeds, further supporting the potential of laser-based stimulation for improving early plant growth responses.^29^ Parveen et al. also evaluated He-Ne laser irradiation in safflower seeds and reported improvements in germination attributes, seed energy metabolism, germination-related enzyme activity, and biochemical metabolites. ^30^ Krawiec et al. investigated He-Ne laser irradiation in scorzonera seeds and observed improvements in germination capacity, radicle length, seedling dry weight, field emergence, and root yield.^31^ Lepossa et al. applied red laser biostimulation to perennial ryegrass seeds and found that, although germination rate was not significantly affected, selected irradiation times improved later shoot growth and biomass-related responses. ^32^ Sevostyanova et al. further reported that red-spectrum laser irradiation can influence germination capacity and early growth processes in mustard and radish seeds. ^33^ Moreover, Li et al. showed that laser diodes can provide precise monochromatic light for plant growth and photosynthetic regulation, with improved growth responses observed in several plant species, including lettuce.^34^ Collectively, these studies indicate that laser irradiation can act as a non-chemical, controllable, and wavelength-selective seed-priming approach. Despite these promising findings, seed responses to laser irradiation strongly depend on plant species, laser wavelength, power density, exposure time, irradiation dose, and growth conditions.

Lettuce *(Lactuca sativa)* is an important leafy vegetable crop and a well-established model system for studying light-mediated seed germination because of its rapid germination, short early growth period, and strong sensitivity to optical stimulation. Unlike many crop seeds, lettuce seeds exhibit a strongly light-dependent germination response. Classical photobiological studies have demonstrated that red and far-red light can reversibly regulate lettuce seed germination through phytochrome-mediated pathways, where red light generally promotes germination and far-red light can suppress or reverse this response. ^35–37^ In addition to red/far-red regulation, blue light in the 400–500 nm spectral region has also been reported to influence lettuce seed germination, although the response may vary depending on dormancy level, cultivar, irradiation conditions, and environmental factors. ^38,39^ These findings indicate that lettuce seeds are responsive to specific wavelengths of light, making them suitable for controlled laser-based seed stimulation. Despite this well-known light sensitivity, most earlier lettuce studies have focused mainly on the fundamental photobiology of germination rather than on the development of laser irradiation as a practical seed-priming technique. One of the few direct laser-based studies on lettuce seeds was reported by Scheuerlein and Braslavsky, who used 15 ns tunable dye-laser flashes in the red spectral range of 620–690 nm to induce germination in photoblastic lettuce seeds.^40^ Their study showed that short laser pulses could trigger a physiological germination response. However, the work was designed primarily to investigate light-induced phytochrome activation rather than to optimize laser irradiation parameters for improving germination performance or early seedling establishment. Although laser biostimulation has been explored as a seed-priming approach in several plant species, its direct application to lettuce seed germination remains comparatively limited. In lettuce, most laser-related studies have focused on post-germination cultivation rather than on direct seed irradiation. For example, Yamazaki et al. used high-power red AlGaInP diode laser lamps at 680 nm for lettuce cultivation and showed that lettuce plants could grow under red diode laser illumination, although biomass production and leaf morphology were not optimal compared with plants grown under conventional lamps.^41^ Similarly, Mori et al. investigated lettuce growth under diode lasers at 440, 660, and 680 nm. They reported that lettuce could grow under laser illumination; however, growth and photosynthetic performance were generally lower than under LED lighting, while vitamin C content increased under laser treatment.^42^ Murase et al. further evaluated a scanning red–green–blue laser projector as an artificial light source for lettuce seedlings and mature plants, focusing on parameters such as plant area, fresh weight, relative growth rate, and plant temperature. ^43^ More recently, combined red/blue laser illumination at 671 and 450 nm has been explored for full-cycle lettuce growth, where physiological and molecular responses were compared with those under white LED illumination.^44^ Together, these studies confirm that lettuce plants can respond to diode laser illumination. However, they mainly address post-germination growth and controlled-environment cultivation rather than laser pretreatment of seeds. This creates an important research gap between the established light sensitivity of lettuce seeds and the practical use of laser irradiation to enhance lettuce germination and early seedling performance. In particular, direct information on CW blue diode-laser pretreatment of lettuce seeds remains limited, especially in the 415–450 nm spectral region. This gap is important because blue light can interact with photoreceptors and early developmental pathways, potentially influencing germination, seedling vigor, and stress adaptation. Moreover, little is known about whether blue laser pretreatment can support lettuce seed germination and seedling establishment under reduced water availability. Therefore, a systematic investigation of laser power, exposure duration, and delivered energy is necessary to identify treatment conditions that can enhance lettuce germination and early seedling growth without inducing inhibitory or stress-related effects.

In the present study, lettuce seeds were treated with a 450 nm CW diode laser to test the hypothesis that an optimized blue-light irradiation dose can regulate early germination dynamics and improve seedling establishment, particularly under reduced water availability. The specific objectives were to (i) determine the effects of laser power, exposure time, and delivered energy on lettuce seed germination; (ii) assess early root and shoot growth as indicators of seedling vigor; and (iii) examine whether blue laser priming can partially compensate for low water supply during germination and early establishment. This work provides a controlled evaluation of blue laser-induced seed stimulation in lettuce and identifies irradiation conditions that may enhance germination, early seedling growth, and early performance under water restriction in a chemical-free and environmentally compatible manner.

## Materials and Methods

The experimental work was carried out in two sequential steps: (i) continuous-wave (CW) laser irradiation of dry lettuce seeds and (ii) germination of the irradiated and non-irradiated seeds under controlled laboratory conditions. The laser irradiation arrangement and the germination setup are schematically presented in Fig. 1(a,b). Dry, non-soaked lettuce seeds were used in all experiments to investigate the direct influence of laser exposure on subsequent germination behavior.

**Figure 1:**
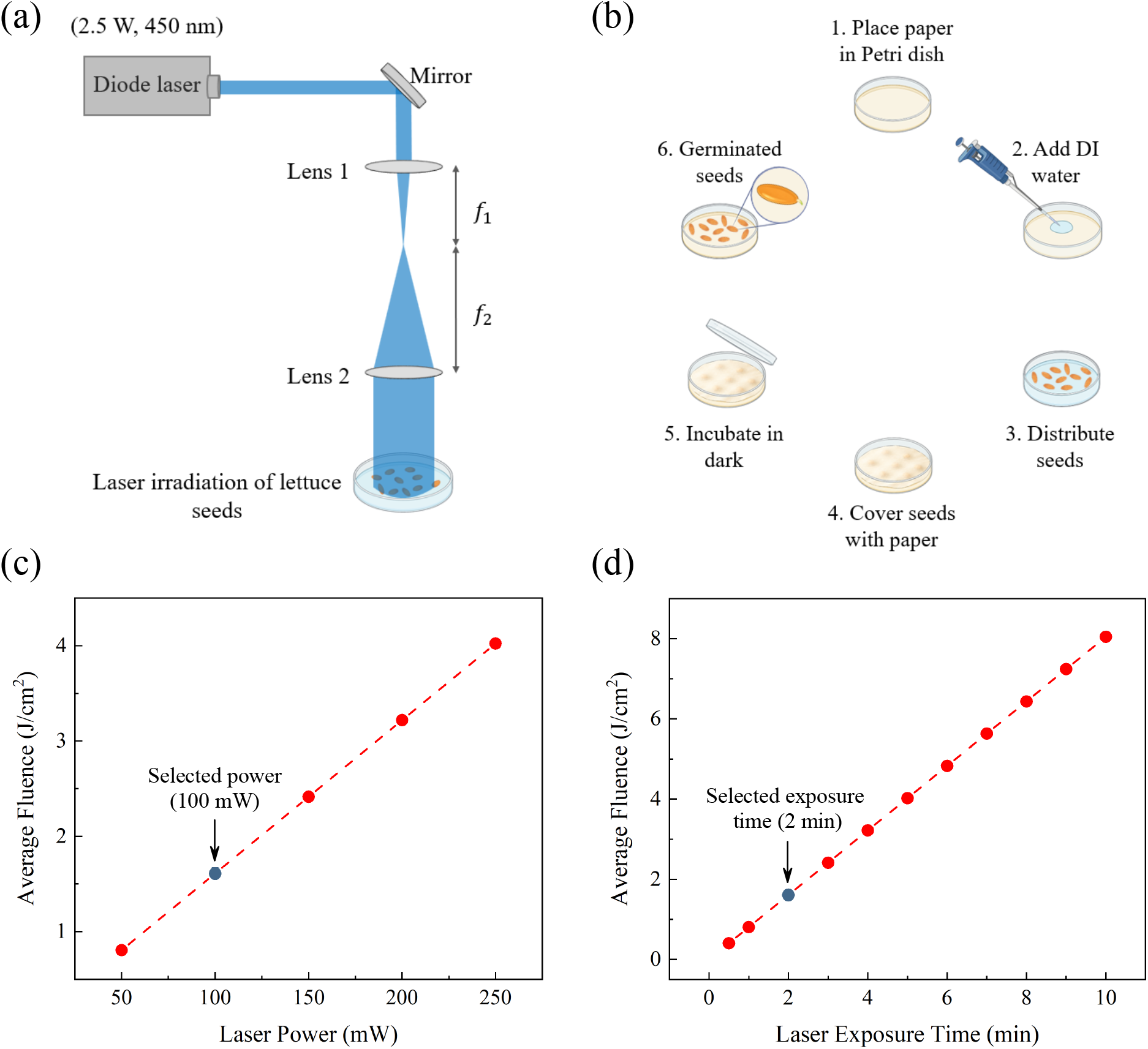
(a) Optical setup for seed irradiation using a 450 nm diode laser with a maximum output power of 2.5 W. The beam was directed by a mirror and expanded through two lenses before irradiating lettuce seeds placed in a Petri dish. (b) Stepwise germination protocol, including placement of tissue paper in the Petri dish, addition of DI water, seed distribution, covering with paper, dark incubation, and evaluation of germinated seeds. Calculated average incident laser fluence as a function of (c) laser power at a fixed exposure time of 2 min and (d) exposure time at a fixed laser power of 100 mW. Fluence was calculated from the measured power at the seed plane and the expanded spot area (*d ≈* 2.5 cm).

### Laser Irradiation Setup and Exposure Procedure

The optical configuration used for seed irradiation is shown in Fig. 1(a). A blue CW diode laser with an emission wavelength of 450 nm and a maximum output power of 2.5 W was employed as the irradiation source. Because the maximum continuous output power of the source was 2.5 W, the system was treated as a Class 4 laser source, and all experiments were performed under controlled laboratory conditions with appropriate laser safety measures. The laser head contained an adjustable lens that modified the beam’s spot size before it entered the external optical system. The laser beam was directed vertically toward the sample plane using a dielectric mirror. After reflection, to increase the exposure area, the beam passed through a two-lens beam-expansion arrangement composed of Lens 1 and Lens 2, with focal lengths of *f*_1_ and *f*_2_, respectively. The expanded beam enabled simultaneous illumination of the seed-containing area inside the Petri dish, reducing spatial intensity variations and minimizing the possibility of localized overheating. The sample position, beam size, and irradiation geometry were carefully adjusted and kept constant for all treatment groups. Before each irradiation experiment, the laser power was measured directly at the seed plane using a calibrated laser power meter. The measurement was performed after the beam had passed through all optical components, including the mirror and lenses, so that optical losses introduced by the setup were included in the measured power. The power-optimization experiments used measured laser powers of 50, 100, 150, 200, and 250 mW at a fixed exposure time of 2 min, whereas the exposure-time experiments used a fixed measured power of 100 mW for durations from 0.5 to 10 min. Each seed group was exposed to only one specific combination of laser power and irradiation duration.

For each experimental sample, 100 dry lettuce seeds were placed in a clean Petri dish and arranged as a single, non-overlapping layer before irradiation. The Petri dish was positioned beneath the expanded laser beam, and the same seed arrangement, sample position, and beam geometry were used for all laser-treated groups. A non-irradiated control group was also prepared using 100 dry lettuce seeds per sample. The control samples were subjected to the same handling and environmental conditions as the treated samples, except that they were not exposed to laser irradiation. Each laser-treatment condition, as well as the control condition, was prepared in three independent replicate samples. Thus, each condition consisted of three Petri dishes, with 100 seeds in each dish. The germination response reported for each condition was calculated as the average of the three independent replicates, and the standard deviation was used to represent experimental uncertainty and generate the corresponding error bars.

To express the irradiation conditions in terms of average incident fluence, the expanded laser spot diameter at the seed plane was 2.4 cm. The corresponding spot area was *A*_spot_ = *π*(*d/*2)^2^ = 4.52 cm^2^. The average projected area of one lettuce seed was approximately 2.78 mm^2^ (0.0278 cm^2^). Therefore, a sample containing 100 seeds had a total projected seed area of approximately 2.78 cm^2^, corresponding to about 61% of the expanded laser spot area. The average incident fluence values shown in Fig. 1(c,d) were calculated as *F* = *Pt/A*_spot_, where *P* is the measured laser power at the seed plane and *t* is the exposure time. For the optimized condition of 100 mW for 2 min, the total incident energy was 12 J and the average incident fluence over the expanded spot was approximately 2.65 J cm*^−^*^2^. These values describe the nominal incident optical dose at the sample plane; they do not account for seed curvature, surface scattering, or seed-to-seed variation in absorbed energy.

### Seed Germination Procedure and Monitoring

Immediately after laser exposure, the treated seeds were transferred to the germination setup shown in Fig. 1(b). Before each germination experiment, all Petri dishes and laboratory materials that came into contact with the seeds were cleaned carefully to reduce the risk of contamination. Sterile disposable Petri dishes were used whenever available. When reusable Petri dishes were used, they were washed thoroughly with neutral laboratory detergent, rinsed repeatedly with deionized water, and dried before use. Ethanol cleaning followed by deionized-water rinsing was also applied where required. Strong chemical cleaning agents, such as aqua regia, were avoided because residual acidic traces or chemical contaminants could affect the germination response. For germination, each Petri dish was first lined with clean tissue paper. A fixed volume of 6 ml of deionized water was added to the tissue paper in each dish for the standard germination experiments. This water volume was kept constant for all control and laser-treated samples in the power- and exposure-time studies to ensure identical hydration conditions. The selected volume was sufficient to moisten the tissue paper uniformly and provide water for seed germination while preventing the formation of a visible pool of free water inside the Petri dish, which could otherwise influence oxygen availability and seedling development. For each sample, 100 lettuce seeds were distributed uniformly over the moistened tissue paper in a single, non-overlapping layer. Adequate spacing was maintained between neighboring seeds to allow clear manual observation of individual seed germination. After seed placement, the seeds were covered with a second sheet of moistened tissue paper to improve moisture contact and reduce dehydration during incubation. The Petri dishes were then covered tightly with their lids to minimize evaporative water loss and maintain a humid environment inside the dishes. The same germination procedure was applied to both the laser-treated samples and the non-irradiated control samples. All Petri dishes were incubated at a controlled temperature of 17–18 *^◦^*C throughout the germination period. The samples were kept in darkness to prevent ambient room light from influencing lettuce seed germination. All handling, inspection, and imaging procedures were performed under dim green LED illumination, which served as a safe-light condition to minimize unintended light-induced effects during monitoring. The control and laser-treated groups were maintained under identical temperature, moisture, light-exposure, and handling conditions, ensuring that the laser irradiation parameters were the only deliberately varied experimental factors. Seed germination was monitored manually during the incubation period after every 4 h. A seed was considered germinated when clear radicle emergence was observed.

### Calculation of Germination Parameters

All germination parameters were calculated independently for each replicate. At each observation time, the cumulative germination percentage *CGP_i_* was calculated as^45^

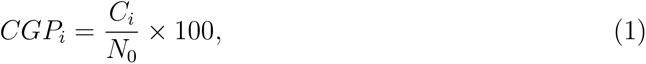

where *N*_0_ is the initial number of seeds and *C_i_* is the cumulative number of seeds germinated by observation time *t_i_*. The cumulative germination count was obtained from 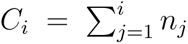, where *n_j_* is the number of newly germinated seeds recorded during the observation interval ending at time *t_j_*. The final germination percentage (FGP) represents the proportion of seeds that had germinated by the end of the 72-h observation period and was calculated as

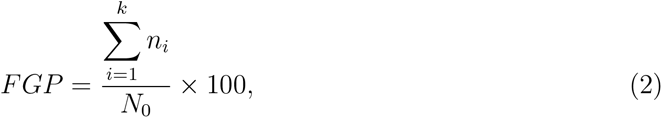

where *k* is the final observation interval.

To describe the average timing of germination, the mean germination time MGT was calculated as the weighted mean of the germination times: ^45,46^

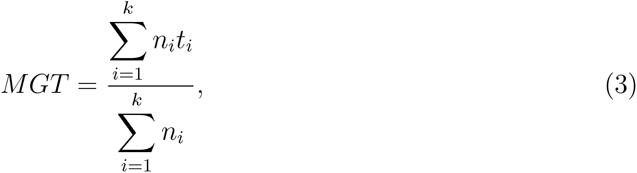

where *n_i_* is the number of seeds that germinated during the interval ending at time *t_i_*. The MGT was expressed in hours, with a lower value indicating that the seeds germinated earlier on average. The mean germination rate (MGR) was calculated as the reciprocal of MGT:^45^

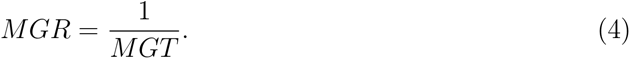

The MGR was expressed in h*^−^*^1^, with higher values indicating faster germination. ^45^ Because MGR was calculated only from the germination times of seeds that successfully germinated, it primarily described the timing of germination and did not directly account for the final germination percentage.

The germination speed index GSI was calculated according to the method of Maguire:^47^

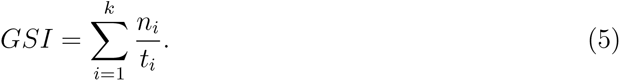

For this calculation, *n_i_*represents the number of newly germinated seeds at each observation interval rather than the cumulative germination count. When *n_i_* was expressed as the number of seeds, GSI had units of seeds h*^−^*^1^. When *n_i_* was expressed as a percentage of the initial seed population, GSI had units of % h*^−^*^1^. A higher GSI indicates earlier germination, a greater number of germinated seeds, or a combination of both. Thus, unlike MGR, GSI reflects both germination timing and germination extent.

The absolute time to 50% germination *T*_50,abs_ was defined as the time required for 50% of the initially sown seeds to germinate. The corresponding cumulative germination threshold was *C^∗^* = 0.5*N*_0_. When this threshold occurred between two consecutive observations, *T*_50,abs_ was estimated by linear interpolation: ^48,49^

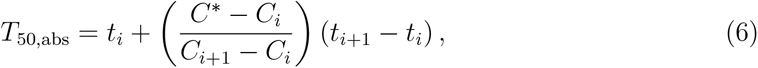

where *C_i_* is the cumulative germination count immediately below the threshold and *C_i_*_+1_ is the cumulative count at or above the threshold, such that *C_i_ < C^∗^ ≤ C_i_*_+1_. The corresponding observation times are denoted by *t_i_* and *t_i_*_+1_. When cumulative germination was expressed as a percentage, the threshold *C^∗^* was replaced by 50%. This interpolation procedure was adapted from methods commonly used to estimate median germination time. ^48^ Unlike the conventional relative *T*_50_, which is defined as the time required to reach 50% of the final number of germinated seeds, *T*_50,abs_ was calculated relative to the initial number of seeds. Therefore, *T*_50,abs_ was considered undefined when the final germination percentage remained below 50%.

Finally, the reproducibility of the laser-treatment response across independent experimental days was evaluated using the relative standard deviation (RSD):^50^

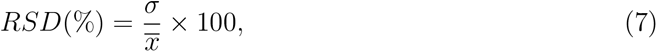

where *σ* is the standard deviation of the final germination percentages measured on different experimental days and *x* is their corresponding mean value for the laser-treated group. A lower RSD indicates less day-to-day variation and, therefore, greater reproducibility of the observed germination response.

### Experimental Design and Data Presentation

The Petri dish was considered the independent experimental unit. Unless otherwise noted, each treatment was represented by three independent Petri-dish replicates, each containing 100 seeds. Germination and seedling-growth parameters were calculated separately for each replicate and are reported as mean values, with standard deviation used to describe variation among replicates in the figures. Treatment responses were interpreted from the magnitude and consistency of the mean responses, the temporal germination profiles, and the doseresponse behavior. Where formal statistical groupings are not shown, terms such as higher, lower, increased, and decreased refer to differences among treatment means rather than to assigned statistical significance.

## Results and Discussion

### Effect of Laser Parameters on Lettuce Seed Germination

Fig. 2 shows the effect of laser power on lettuce seed germination at a fixed irradiation time of 2 min. Each treatment consisted of three independent replicates, with 100 seeds per replicate. Non-irradiated seeds were used as the control, while the remaining groups were irradiated at laser powers of 50, 100, 150, 200, or 250 mW. All groups were maintained under identical germination conditions. As shown in Fig. 2(b), no radicle emergence was observed during the first 12 h in either the control or laser-treated groups. Germination then increased rapidly, particularly between approximately 16 and 36 h, before approaching a plateau after about 40 h. The cumulative germination curves of the laser-treated groups were steeper than that of the control, indicating that more seeds germinated during the main germination phase. The control group reached a final germination percentage of approximately 68%, whereas the laser-treated groups reached approximately 81–92%. The largest improvement occurred between the control and the 100 mW treatment. In comparison, the final germination percentages of the 100–250 mW groups varied within a relatively narrow range, suggesting that the biostimulatory response began to approach saturation at approximately 100 mW. However, because the error ranges of several laser-treated groups overlap, these results should not be interpreted as a strictly monotonic power-dependent response without further statistical confirmation. The MGR increased moderately from approximately 0.040 h*^−^*^1^ in the control to 0.042–0.045 h*^−^*^1^ in the laser-treated groups, as shown in Fig. 2(c). This moderate increase in MGR is consistent with the steeper cumulative germination curves because MGR describes the average timing of germinated seeds, whereas the cumulative curve and final germination percentage also reflect how many seeds germinated. Laser irradiation therefore increased the number of seeds germinating during the main germination phase but produced a smaller reduction in their average germination time. In addition, the germination of additional seeds at later observation times may increase the weighted mean germination time and partly offset the effect of earlier germination. These results indicate that laser treatment primarily improved germination capacity, with a smaller improvement in the average germination rate. A clearer response was observed in the GSI, which increased from approximately 3.0 in the control to 3.8–4.4 in the irradiated groups, as shown in Fig. 2(d). The GSI was calculated using Eq. (5). Unlike MGR, GSI is influenced by both the number of seeds that germinate and the timing of their germination. The higher GSI values therefore indicate that laser irradiation increased the number of seeds germinating during the early and intermediate observation periods. This trend agrees with the final germination results presented in Fig. 2(e), where all irradiated groups exhibited higher mean final germination percentages than the control. Although the highest mean final germination percentage was recorded at 250 mW, the differences among the 100–250 mW treatments were relatively small. Thus, a substantial proportion of the maximum germination response had already been achieved at 100 mW. Taking into account the small additional improvement obtained at higher powers, 100 mW was selected as a suitable power for subsequent exposure-time experiments. Representative images of three-day-old seedlings from the control, 50 mW, and 100 mW groups are shown in Fig. 2(f)–(h), respectively. Seedlings developed from irradiated seeds, particularly those treated at 100 mW, showed greater radicle elongation and more advanced early development than the control seedlings. Although these images provide qualitative evidence, they support the improvements observed in the calculated germination parameters.

**Figure 2:**
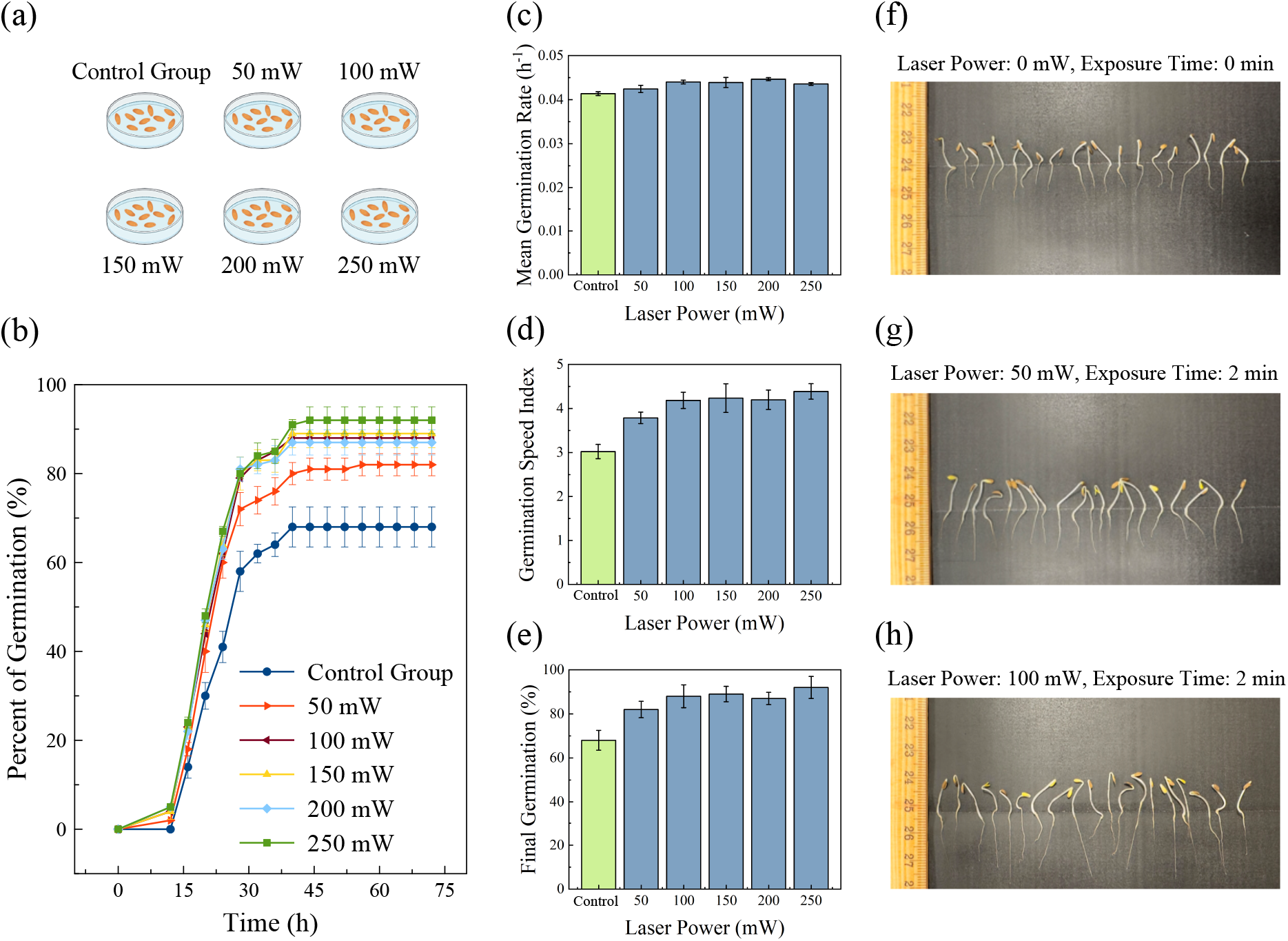
(a) Schematic representation of the control group and lettuce seed groups irradiated with laser powers of 50, 100, 150, 200, and 250 mW for a fixed exposure time of 2 min. (b) Time-dependent germination percentage of the control and laser-treated groups recorded over 72 h. (c-e) Mean germination rate, germination speed index, and final germination percentage, respectively, as functions of laser power. Error bars represent the standard deviation among three independent Petri-dish replicates. (f-h) Representative digital images of three-day-old lettuce seedlings from the control group (0 mW, 0 min), the 50 mW-treated group, and the 100 mW-treated group, respectively.

Following the power-dependent study, the effect of irradiation duration was investigated at a fixed laser power of 100 mW, as shown in Fig. 3. Non-irradiated seeds were used as the control, while the treatment groups were exposed to the laser for durations ranging from 0.5 to 10 min. Each condition consisted of three independent replicates, with 100 seeds per replicate. Because the laser power remained constant, increasing the exposure time progressively increased the incident optical energy delivered to the seeds. The cumulative germination percentage (CGP), calculated as described in the preceding subsection, is presented in Fig. 3(b). No radicle emergence was observed during the first 12 h. Germination then increased rapidly between approximately 16 and 36 h before approaching a plateau after about 40 h. Short irradiation durations increased both the rate at which germinated seeds accumulated and the final percentage of germination compared with the control. Among the tested conditions, the 2 min treatment produced the steepest CGP curve and the highest mean final germination percentage of approximately 93%, compared with approximately 67% for the control. Irradiation for 0.5 and 1 min also improved germination, although the responses were smaller than that obtained at 2 min. This result indicates that short exposures were sufficient to initiate a stimulatory response, but an exposure time of 2 min was required to produce the maximum response under the present experimental conditions. Increasing the irradiation duration beyond 2 min progressively reduced the beneficial effect. The final germination percentage decreased from approximately 93% at 2 min to approximately 87% at 3 min and continued to decline at longer exposure times. After 10 min of irradiation, the final germination percentage decreased to approximately 64%, which was slightly below the control value. A similar non-linear response to irradiation duration has been reported for mung bean seeds treated with CW lasers at 488 and 632.8 nm; irradiation for 2 min produced the greatest seedling development, whereas the response decreased after exposures of 5 and 10 min.^51^ Higher laser doses have also been reported to inhibit soybean-seed germination after beneficial responses at shorter exposure durations. ^52^ These findings support the existence of an optimum irradiation range within which laser treatment is beneficial. However, the position of this optimum depends on several factors, including plant species, wavelength, irradiance, seed condition, and experimental environment.^26,52–55^ The MGR presented in Fig. 3(c) showed a similar exposure-time-dependent response. The control group had an MGR of approximately 0.040 h*^−^*^1^. The value decreased slightly after 0.5 min of irradiation and then reached a maximum of approximately 0.045 h*^−^*^1^ at 2 min. The slightly lower MGR after 0.5 min, despite a higher final germination percentage than the control, can be explained by the different information represented by these two parameters. The final germination percentage describes the proportion of the initial seed population that successfully germinated, whereas MGR reflects the average germination timing of only those seeds that germinated. Thus, the 0.5 min treatment allowed more seeds to germinate but did not reduce their average germination time. Beyond 2 min, MGR decreased progressively and reached approximately 0.034 h*^−^*^1^ at 10 min, indicating that prolonged irradiation delayed the average timing of radicle emergence. A closely corresponding trend was observed for GSI, as shown in Fig. 3(d). GSI increased from approximately 2.8 in the control to a maximum of approximately 4.5 after 2 min of irradiation and then declined with increasing exposure time. Because GSI accounts for both the number of germinated seeds and their germination timing, its maximum at 2 min confirms that this treatment produced the best overall germination performance among the investigated conditions. The decrease in GSI at longer exposure times indicates that prolonged irradiation reduced the number of successfully germinated seeds, delayed germination, or produced both effects. This trend agrees with the final germination results shown in Fig. 3(e), which improved up to 2 min and then progressively declined, eventually falling below the control after 10 min. The decline observed at longer exposure times may indicate a transition from beneficial photobiostimulation to irradiation-induced stress. ^56,57^ At an appropriate optical dose, laser treatment can stimulate physiological processes associated with seed germination and early seedling development.^22,58^ In contrast, excessive irradiation may disturb normal cellular processes and consequently reduce germination and seedling growth.^55–57,59^ However, the physiological and biochemical processes involved were not measured in the present study. Therefore, the mechanism responsible for the reduced response at longer exposure times cannot be determined conclusively and requires further investigation. The representative images of three-day-old seedlings shown in Fig. 3(f)–(h) provide qualitative support for the quantitative germination results. Seedlings developed from seeds irradiated for 2 min showed more advanced radicle development than the control seedlings, whereas those from the 10 min treatment exhibited visibly restricted development. These images suggest that the effect of irradiation duration continued beyond radicle emergence and influenced early post-germination growth. However, conclusions regarding root and shoot enhancement should be based primarily on quantitative length measurements rather than representative images alone. Overall, the results identify 2 min at 100 mW as the most effective irradiation condition for improving lettuce seed germination among the exposure durations investigated.

**Figure 3:**
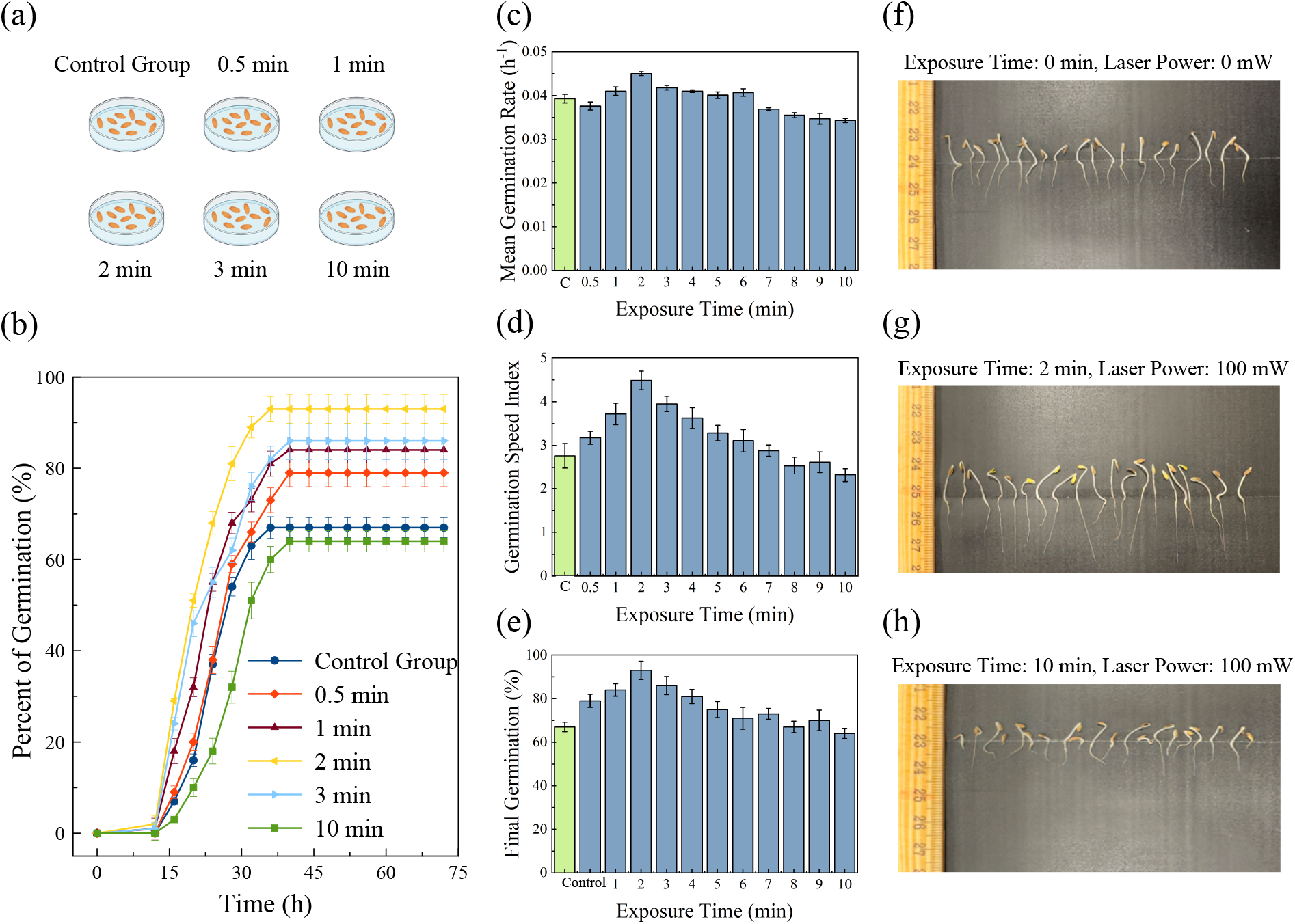
(a) Schematic representation of the control group and lettuce seed groups irradiated at a fixed laser power of 100 mW for different exposure times. (b) Time-dependent germination percentage of the control and laser-treated groups recorded over 72 h. (c-e) Mean germination rate, germination speed index, and final germination percentage, respectively, as functions of laser exposure time. Error bars represent the standard deviation among three independent Petri-dish replicates. (f-h) Representative digital images of three-day-old lettuce seedlings from the control group (0 min, 0 mW), the 2 min-treated group, and the 10 min-treated group, respectively.

Fig. 4(a) schematically illustrates the transition from seed germination to early seedling establishment. Germination begins with imbibition, during which water uptake causes seed expansion and initiates the physiological processes required for embryo growth. Germination is completed by the emergence of the radicle through the seed covering, which subsequently develops into the primary root. With continued growth, the hypocotyl elongates and the young shoot begins to emerge from the seed coat.^60^ At the same time, the root system extends and develops additional lateral roots, while the cotyledons expand above the emerging shoot. As development proceeds, the seed coat is gradually shed, and both the shoot and root systems continue to elongate, producing an established lettuce seedling.^61,62^ Root and shoot lengths were therefore measured as indicators of early seedling growth and vigor rather than germination alone. ^63^ Fig. 4(b,c) show the effects of laser power on the mean shoot and root lengths, respectively, at a fixed irradiation time of 2 min. The control group showed the lowest shoot length, indicating limited early shoot development in non-irradiated seeds. In comparison, all laser-treated groups showed improved shoot elongation. Shoot length increased noticeably at 50 mW and reached its maximum value around 100 mW. At higher laser powers, shoot length remained higher than the control, although the variation among 150, 200, and 250 mW was relatively small. This pattern suggests that laser irradiation promoted shoot development, but the shoot response became less sensitive once a sufficient stimulation level was reached.

**Figure 4:**
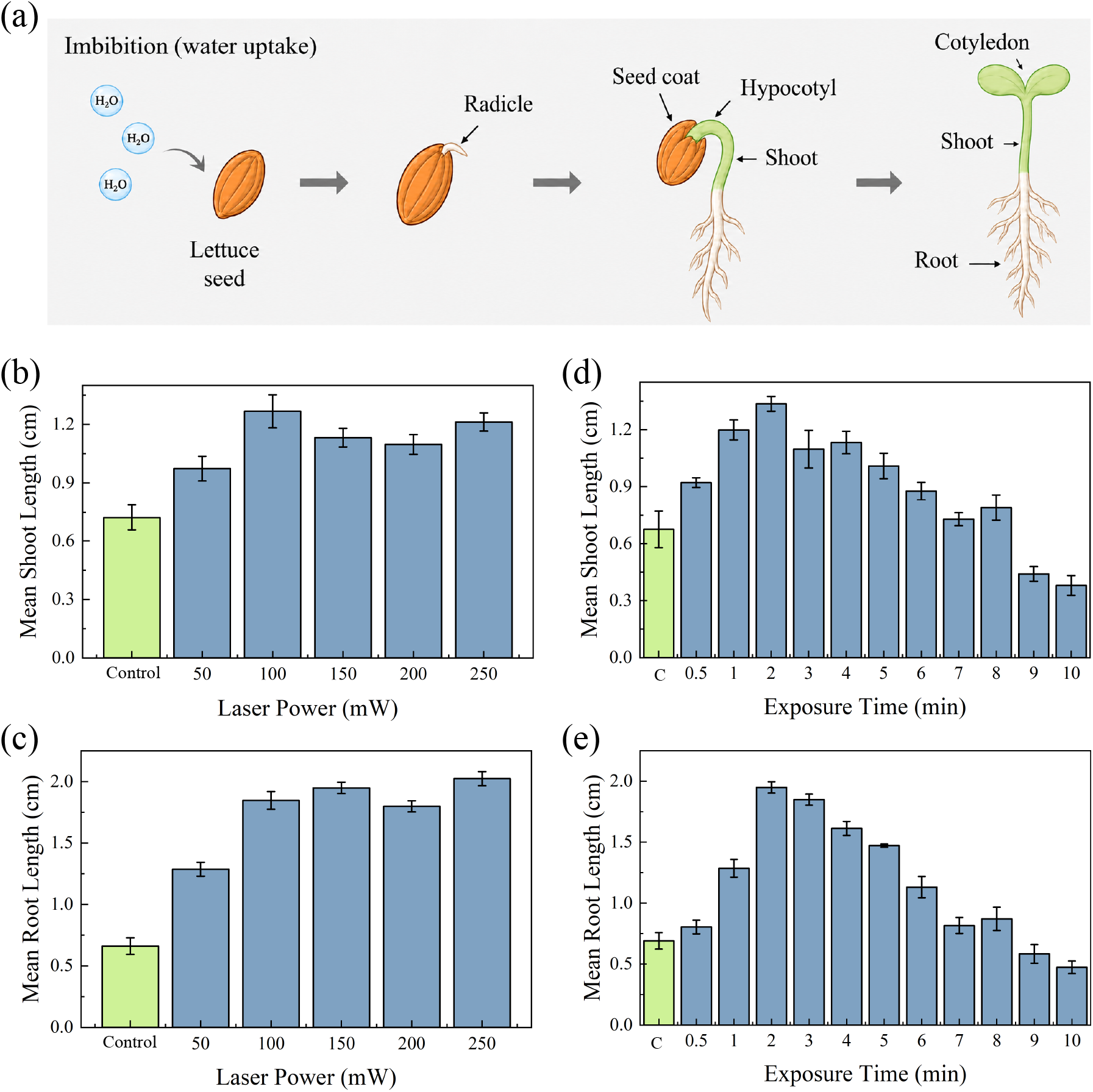
(a) Schematic representation of lettuce seed germination, showing water uptake during imbibition, radicle emergence, seed-coat rupture, hypocotyl development, shoot formation, and root growth. (b,c) Mean shoot length and mean root length of lettuce seedlings as functions of laser power. (d,e) Mean shoot length and mean root length as functions of laser exposure time. The control group represents non-irradiated seeds, while all treated groups were irradiated under controlled conditions with the water volume kept constant at 6 ml. Error bars represent the standard deviation among three independent Petri-dish replicates.

A stronger effect was observed in the root-length data shown in Fig. 4(c). The control seedlings had the shortest roots, while laser-treated seedlings showed a clear increase in root elongation. The increase in root length was more pronounced than the increase in shoot length, indicating that laser treatment had a stronger influence on early root development. This response is biologically plausible because the radicle is the first organ to emerge from the seed and root growth responds rapidly to changes in hydration, metabolic activation, and seed vigor. Laser irradiation may enhance water uptake or activate early physiological processes that support radicle emergence and root elongation. In addition, shoot elongation and cotyledon expansion occur slightly later and depend more strongly on reserve mobilization and hypocotyl development. Therefore, even when laser treatment improves overall seedling vigor, root length may show a larger and earlier response than shoot length.

Fig. 4(d,e) show the effect of exposure time on mean shoot length and mean root length at a fixed laser power of 100 mW. Shoot length increased with exposure time up to 2 min, where the maximum value was observed. When exposure time was increased beyond 2 min, shoot length gradually decreased, with a strong reduction at 9 and 10 min. A similar but more pronounced trend was observed for root length in Fig. 4(e). Root length increased strongly from the control to the 1 and 2 min treatments, with maximum root elongation obtained at 2 min. After this optimum exposure time, root length decreased progressively with increasing irradiation time. The 10 min-treated seedlings showed much shorter roots than the 2 min-treated seedlings, confirming that excessive laser exposure suppressed early root growth. This result is consistent with the germination data, where 2 min exposure produced the best germination response, while longer exposure times reduced overall germination performance. The decrease in both shoot and root length at longer exposure times suggests that the laser effect is dose-dependent. At an appropriate dose, laser irradiation can act as a biostimulatory treatment by supporting water absorption, metabolic activation, and early cell division and elongation. However, excessive delivered energy may induce stress in seed tissues.^64^ Prolonged irradiation may cause photothermal or photochemical stress, disturb cellular membranes, or increase oxidative stress, all of which can reduce seedling vigor. Root growth is especially sensitive to such stress because the root tip contains actively dividing and elongating cells. Therefore, excessive exposure can strongly inhibit root elongation and lead to weaker seedling development.

To determine whether the observed improvement was associated with controlled laser irradiation, the germination response of laser-treated seeds was compared with that of sunlight-exposed seeds, as shown in Fig. 5. Three experimental groups were investigated: a non-irradiated control group, a laser-treated group, and a sunlight-treated group. The laser- and sunlight-treated seeds were exposed for the same duration of 2 min, while the control seeds received no light treatment before germination. All groups were subsequently maintained under identical germination conditions. The cumulative germination profiles are presented in Fig. 5(a). The control group reached a final germination percentage of approximately 73%. Sunlight exposure produced only a small improvement, with the final germination percentage remaining slightly above that of the control. In contrast, the laser-treated group reached a substantially higher final germination percentage of approximately 93%. The laser-treated seeds also germinated more rapidly during the main germination phase, as indicated by the steeper increase in cumulative germination between almost 16 and 32 h. These results demonstrate that, under the exposure conditions used in this study, the 450 nm laser treatment produced a stronger germination response than either sunlight exposure or no light treatment. The different responses to laser and sunlight exposure may be related to differences in their spectral and dosimetric characteristics. Sunlight contains a broad range of ultraviolet, visible, and infrared wavelengths, and its spectral irradiance can vary with the time of day, weather, atmospheric conditions, and experimental location. In contrast, the laser delivered light at a defined wavelength, power, and exposure time. The 450 nm irradiation therefore provided a controlled and reproducible blue-light stimulus. Although sunlight also contains blue light, the spectral photon dose received by the seeds near 450 nm was not measured or matched to that of the laser. Consequently, the stronger response of the laser-treated group may be associated with its controlled wavelength and optical dose, but the present comparison does not establish that monochromatic laser light is inherently more effective than sunlight under all conditions. The possible biological basis of the observed response is illustrated schematically in Fig. 5(b). Plants contain several families of photoreceptors that respond to different regions of the optical spectrum.^20,64,65^ Phytochromes primarily perceive red and far-red light, whereas cryptochromes and phototropins are sensitive mainly to blue and UV-A light. These photoreceptors allow plants to distinguish differences in light wavelength and initiate appropriate physiological responses. The classical stimulation of lettuce seed germination by red light, for example, is mainly associated with phytochrome activation.^11,35^ Because the laser used in the present study emitted at 450 nm, cryptochromes, particularly CRY1 and CRY2, represent possible photoreceptors involved in the response.^64,65^ As proposed in Fig. 5(b), absorption of blue photons by cryptochromes may alter the redox state of their flavin chromophore and generate an active signaling state.^11,20^ This photochemical event can initiate downstream light-signaling processes that regulate plant growth and development. In germinating seeds, light-dependent signaling may interact with the hormonal balance between gibberellins (GA) and abscisic acid (ABA).^66,67^ In general, GA promotes processes associated with reserve mobilization, embryo growth, and radicle emergence, whereas ABA helps maintain seed dormancy and opposes germination.^68,69^ Previous studies have shown that light and photoreceptor signaling can influence GA and ABA metabolism during seed germination.^20,64^ Based on these established responses, the enhanced germination observed after blue-laser treatment may involve blue-light perception by cryptochromes, followed by redox-associated signaling and possible modulation of the GA/ABA balance. Such changes could support metabolic activation, reserve mobilization, embryonic growth, and eventual radicle emergence.^65^ However, Fig. 5(b) represents only a simplified, literature-based conceptual pathway. Cryptochrome activation, cellular redox status, and GA and ABA concentrations were not measured in the present study. Therefore, the proposed pathway should not be interpreted as a mechanism experimentally confirmed by the current results.^20,64^ The comparison between Figs. 5(a) and 5(b) therefore connects the measured germination response with a possible wavelength-sensitive biological explanation. Fig. 5(a) experimentally demonstrates that controlled 450 nm irradiation produced a stronger response than short-duration sunlight exposure, while Fig. 5(b) illustrates how blue-light perception could contribute to this response. Although longer sunlight exposure might produce a different result, this possibility was not investigated and cannot be concluded from the present experiment. The main advantage demonstrated here is that laser irradiation provides a controllable and reproducible optical treatment in which the wavelength, power, and exposure time can be accurately adjusted. ^70^

**Figure 5:**
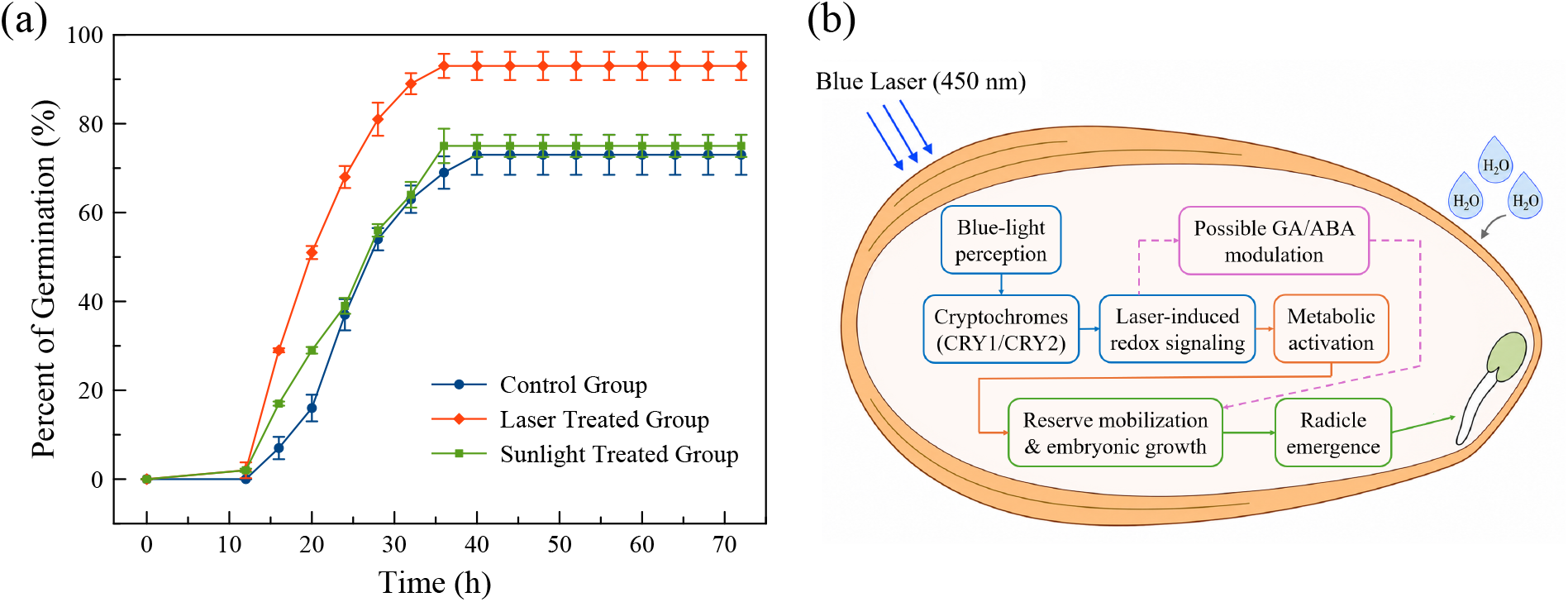
(a) Germination percentage of lettuce seeds over time under different treatment conditions: control, laser-treated, and sunlight-treated groups. All groups were germinated under the same conditions. (b) Simplified literature-based pathway illustrating the proposed role of cryptochrome-mediated blue-light perception in lettuce seed germination.

To evaluate the reproducibility and stability of the optimized laser treatment, the germination response of lettuce seeds treated at 100 mW for 2 min was monitored over 22 different experimental days under the same water condition of 6 ml, as shown in Fig. 6. The x-axis represents independent day-to-day experimental runs, not seedling age. The purpose of this experiment was to determine whether the selected laser condition could consistently improve germination over repeated trials. The RSD was calculated to estimate the day-to-day variation in germination percentage using Eq. (7). As shown in Fig. 6, the laser-treated group showed a relatively high germination percentage during most of the experimental days. In the initial days, germination remained close to 90–95%, confirming the strong positive effect of the optimized laser treatment. Although some day-to-day fluctuation was observed, most values remained above or close to the average germination level. The calculated RSD value of 8.65% indicates that the laser treatment produced a reasonably reproducible germination response over repeated experiments. However, a gradual decline in germination percentage was observed during the later experimental days, and by day 22, the response of the laser-treated group approached that of the control. A previous study reported that laser-induced changes in germination-related hormonal and gene-expression responses were time-dependent. The differences in GA and ABA levels and in the GA/ABA ratio between the irradiated and control groups were more pronounced during the earlier stages after irradiation but gradually became smaller, eventually approaching the corresponding control values at later stages. ^58^ Similarly, the expression of genes associated with germination, hormonal regulation, and light-responsive signaling varied over time, with several initially stimulated responses declining or changing during the later stages.^58^ These findings suggest that laser-induced biostimulatory processes are dynamic and may gradually weaken with increasing time after irradiation. Such a time-dependent reduction in the laser-induced response may help explain why the germination advantage observed in the present study gradually decreased and approached the control level during the later experimental days. However, GA, ABA, and other related biochemical or gene-expression responses were not measured in the present study. Therefore, this explanation remains a literature-based interpretation and cannot be considered a mechanism experimentally confirmed for the lettuce seeds investigated here. Changes in seed physiological condition during storage, together with natural seed-to-seed differences and minor environmental variations, may also have contributed to the observed decline. Consequently, the laser-treated group maintained improved germination over most of the 22-day period, demonstrating that irradiation at 100 mW for 2 min produced a generally repeatable biostimulatory effect under the tested conditions.

**Figure 6:**
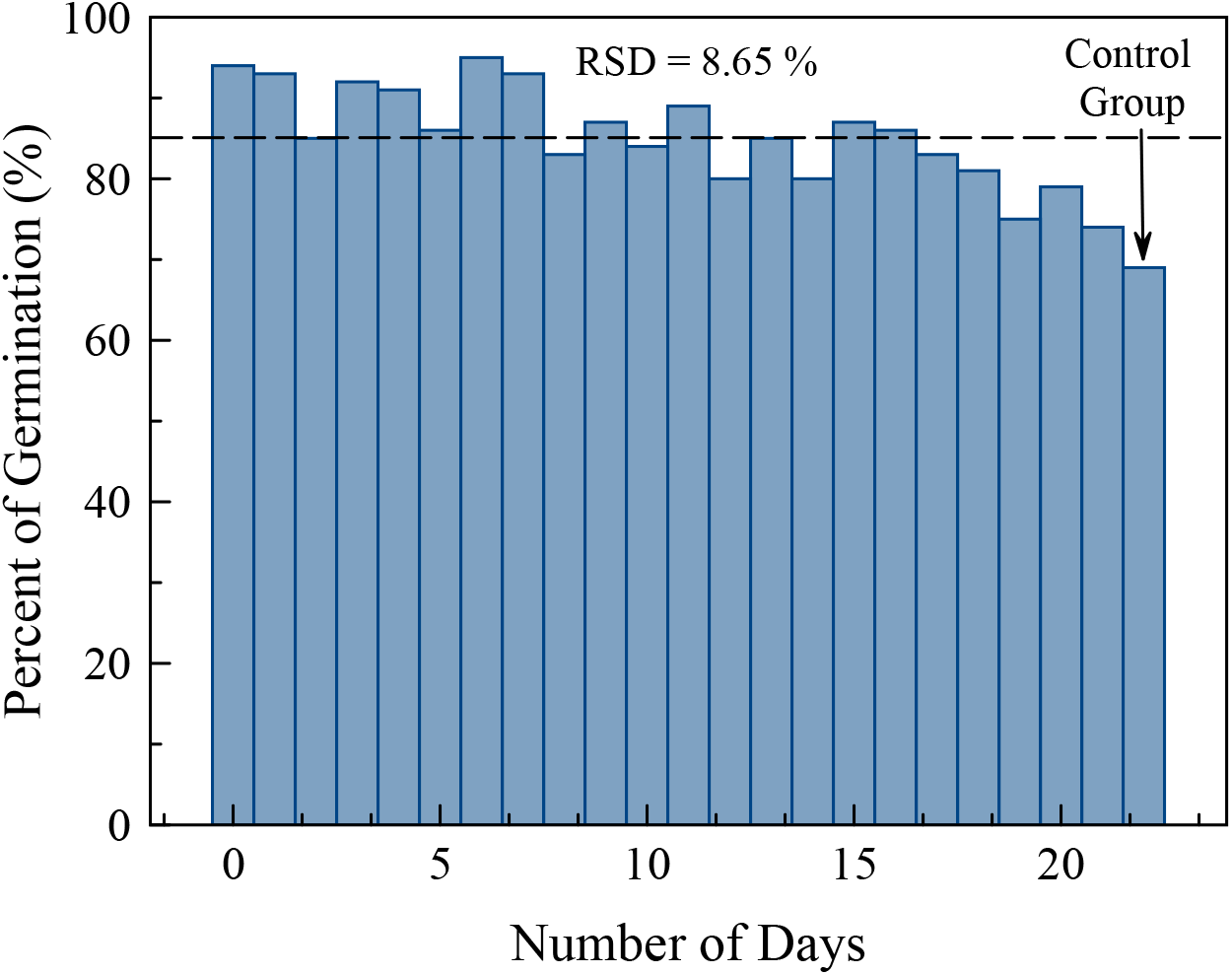
Day-to-day variation in the germination percentage of the laser-treated group exposed for 2 min at 100 mW under 6 ml water conditions over 22 experimental days. The bars represent the germination percentage recorded on each day, while the dashed horizontal line indicates the average germination level of the treated group. The calculated relative standard deviation (RSD = 8.65%) reflects the variation in germination over time. By the 22nd day, the germination percentage of the treated group became comparable to that of the control group.

### Effect of Reduced Water Availability on Lettuce Seed Germination and Early Seedling Growth

The ability of laser pretreatment to improve lettuce-seed performance under water-limited conditions was evaluated by progressively reducing the volume of deionized water supplied to each Petri dish from the standard volume of 6 ml to 4, 2, 1, and 0.5 ml. This assay should be interpreted as reduced water availability during Petri-dish germination rather than as a complete drought-stress model, because water limitation was imposed by changing the supplied volume rather than by controlling soil matric potential or osmotic potential. For each treatment, the specified water volume was distributed as uniformly as possible over the supporting tissue to ensure that all seeds had comparable access to moisture. Particular care was taken at the two lowest water volumes, 0.5 and 1 ml, to prevent water from becoming localized within a limited region of the Petri dish. Therefore, the observed differences primarily reflect changes in the total amount of water available for imbibition and subsequent seedling development rather than differences in water distribution. At each water level, the seeds were divided into a non-irradiated control group and a laser-treated group. The treated seeds were irradiated at 100 mW for 2 min, corresponding to the optimized irradiation condition identified in the preceding experiments, whereas the control seeds were handled identically but received no laser exposure. Three independent replicates, each containing 100 seeds, were examined for every combination of treatment and water volume. Accordingly, the values presented in Fig. 7 represent the mean germination percentage, and the error bars indicate the standard deviation among the three replicates.

**Figure 7:**
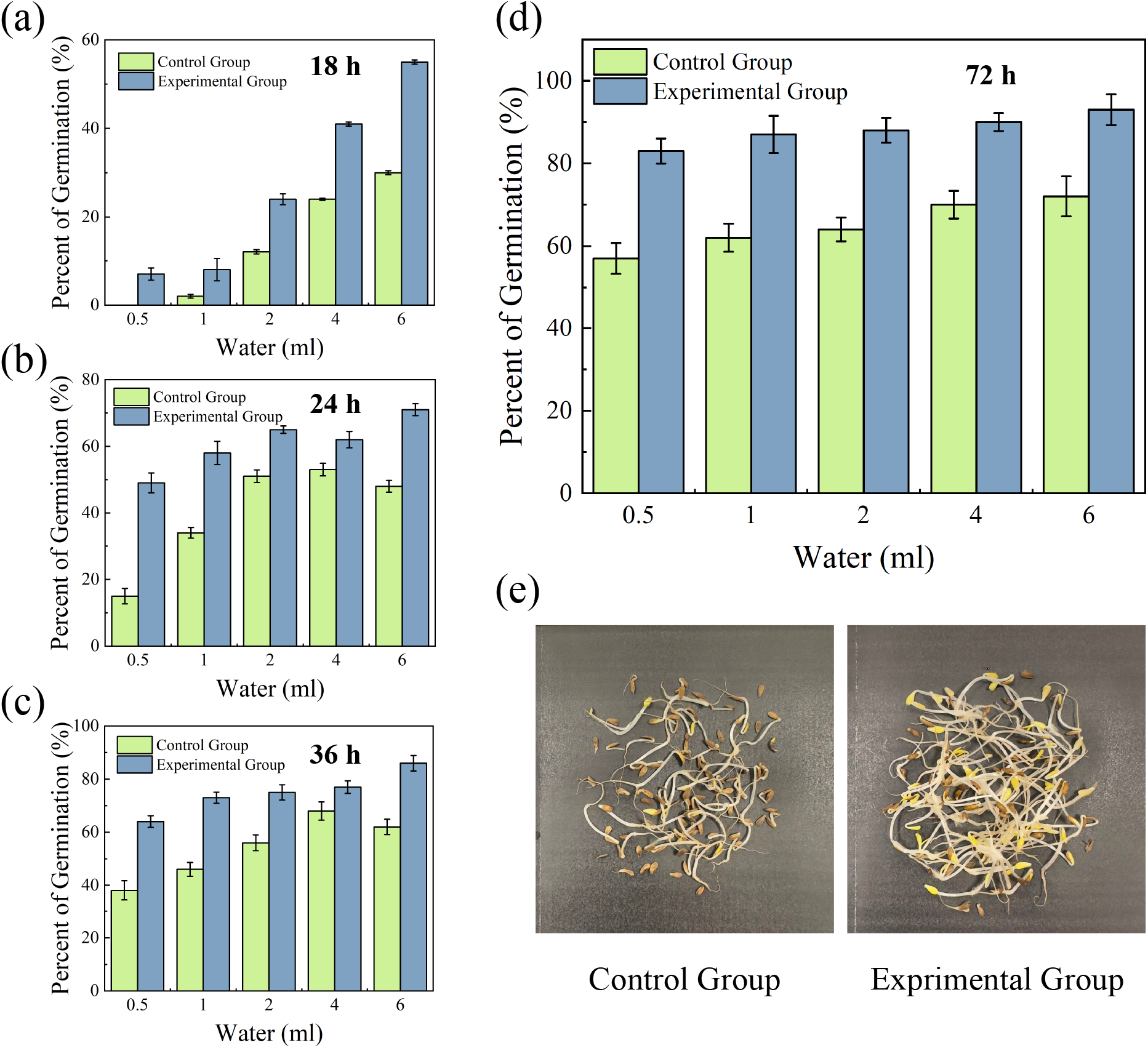
Effect of reduced water availability on the germination and early post-germination growth of lettuce seeds. Germination percentages of non-irradiated control seeds and laser-treated seeds supplied with 0.5, 1, 2, 4, or 6 ml of deionized water were recorded after (a) 18 h, (b) 24 h, (c) 36 h, and (d) 72 h. Laser-treated seeds were irradiated at 100 mW for 2 min before germination, whereas the control seeds received no laser exposure. Each condition consisted of three independent replicates, with 100 seeds per replicate. Error bars represent the standard deviation among the three independent replicates. (e) Representative images of the control and laser-treated seedlings grown with 1 ml of water, showing improved root and shoot development in the laser-treated group.

The early germination response recorded after 18 h is presented in Fig. 7(a). At this stage, germination was strongly influenced by both water availability and laser pretreatment. The control seeds exhibited little or almost no germination at the two lowest water volumes of 0.5 and 1 ml, whereas their germination percentage gradually increased as the supplied water volume was raised from 2 to 6 ml. This behavior is consistent with the essential role of water during seed imbibition. Adequate hydration is required for seed-coat softening, membrane reorganization, enzyme activation, reserve mobilization, and the resumption of embryo growth.^71–73^ Consequently, restricting the available water can delay these processes and postpone radicle emergence.^73–75^ In contrast, the laser-treated seeds exhibited greater germination than the corresponding control seeds at every tested water volume after 18 h. This improvement was observed not only under the well-watered 6 ml condition but also under the strongly water-limited conditions of 0.5 and 1 ml. For example, while the control group showed negligible germination at 0.5 ml, radicle emergence had already begun in the laser-treated group. Clear differences between the two groups were also observed at 2, 4, and 6 ml. These findings indicate that laser pretreatment accelerated the onset of germination across the entire range of water availability. The treatment did not remove the dependence of germination on water, because germination still generally increased with increasing water volume. However, it enabled a larger proportion of seeds to complete the initial germination processes within the same period, indicating partial compensation for low water supply during early imbibition and radicle emergence.

The results obtained after 24 h, shown in Fig. 7(b), further demonstrate the faster germination kinetics of the laser-treated seeds. Germination increased in both groups as additional time became available for water uptake and metabolic reactivation. However, the laser-treated group maintained a higher mean germination percentage than the corresponding control group at all five water volumes. The difference remained particularly apparent at 0.5 and 1 ml, indicating that the laser-induced response was not restricted to seeds supplied with abundant water. Even under limited moisture availability, the treated seeds initiated germination earlier and in greater numbers than the non-irradiated seeds. A similar response was observed after 36 h [Fig. 7(c)]. Although the germination percentages of both groups continued to increase, the laser-treated seeds remained ahead of the control seeds across the complete water-volume range. The persistence of this response from 18 to 36 h demonstrates that the observed enhancement was not confined to a single observation time. Instead, laser pretreatment produced a sustained acceleration of germination during the period of greatest germination activity. Therefore, its principal early effect was to improve germination speed under both sufficient and restricted water availability.

The final germination percentages measured after 72 h are presented in Fig. 7(d). By this time, germination had approached a plateau, with little or no additional germination observed thereafter. The dependence of germination on water volume was less pronounced at 72 h than during the earlier observation periods, particularly for the laser-treated group. Although the control seeds also achieved considerable final germination under water-limited conditions, their mean germination percentages remained lower than those of the laser-treated seeds at every water volume. The control group increased from approximately 57% at 0.5 ml to 72% at 6 ml, whereas the laser-treated group increased from approximately 83% to 93% over the same water range. Thus, reducing the available water caused a moderate decline in final germination, particularly in the control group, whereas laser pretreatment partially mitigated this decline. These results also suggest that water limitation delayed germination more strongly than it completely prevented germination in many viable seeds. However, statements regarding statistical significance should be made only after applying the appropriate statistical comparisons. The germination and seedling-growth parameters summarized in Table 1 provide additional quantitative support for the responses observed in Fig. 7. The differences between the laser-treated and control groups were particularly evident under the most severe water limitation. At 0.5 ml, the laser-treated group achieved a final germination percentage of 83%, compared with 57% for the control group. Under the same condition, laser treatment reduced the MGT from 33.60 to 28.46 h and the absolute time required to reach 50% germination (*T*_50,abs_) from 44.89 to 25.00 h. The treated seeds also exhibited a higher MGR, increasing from 0.0298 to 0.0351 h*^−^*^1^, and a higher GSI, increasing from 1.83 to 3.27. Similar improvements were observed at 1 ml, where *T*_50,abs_ decreased from 38.29 h in the control group to 22.00 h in the laser-treated group. These reductions in MGT and *T*_50,abs_, together with the generally higher GSI values, confirm that laser pretreatment primarily promoted faster and more synchronized germination, especially when water availability was strongly restricted. An important distinction must, however, be made between seed germination and subsequent seedling development. Germination percentage is commonly determined on the basis of visible radicle emergence and therefore indicates successful completion of the initial germination process.

**Table 1:** Germination and early seedling-growth parameters of lettuce seeds under different water-availability conditions for the laser-treated and control groups. Values represent mean responses from three independent Petri-dish replicates, each containing 100 seeds.

| Group | Water level (ml) | Final germination (%) | MGT (h) | MGR ($\text{h}^{-1}$) | GSI | $T_{50,abs}$ (h) | Mean shoot length (cm) | Mean root length (cm) |
| --- | --- | --- | --- | --- | --- | --- | --- | --- |
| **Laser-treated** | 6 | 93.0 | 22.90 | 0.0437 | 4.44 | 17.62 | 1.495 | 1.267 |
|  | 4 | 90.0 | 24.60 | 0.0407 | 4.06 | 19.20 | 1.402 | 1.335 |
|  | 2 | 88.0 | 24.91 | 0.0402 | 3.87 | 19.93 | 1.335 | 1.212 |
|  | 1 | 87.0 | 26.85 | 0.0372 | 3.57 | 22.00 | 1.397 | 1.289 |
|  | 0.5 | 83.0 | 28.46 | 0.0351 | 3.27 | 25.00 | 1.354 | 1.165 |
| **Control** | 6 | 72.0 | 24.94 | 0.0401 | 3.18 | 25.33 | 0.675 | 0.653 |
|  | 4 | 70.0 | 23.94 | 0.0418 | 3.11 | 23.43 | 0.522 | 0.492 |
|  | 2 | 64.0 | 25.00 | 0.0400 | 2.76 | 23.83 | 0.354 | 0.313 |
|  | 1 | 62.0 | 29.20 | 0.0342 | 2.37 | 38.29 | 0.221 | 0.198 |
|  | 0.5 | 57.0 | 33.60 | 0.0298 | 1.83 | 44.89 | 0.180 | 0.135 |

However, this criterion does not necessarily demonstrate that the resulting seedling can maintain root and shoot growth under continued water limitation. This distinction is illustrated by the representative images in Fig. 7(e), which compare control and laser-treated seeds supplied with 1 ml of water. Although a considerable proportion of the control seeds eventually germinated, their post-germination development remained limited. The resulting seedlings generally exhibited short and poorly developed roots and shoots, suggesting that the available water was sufficient to initiate radicle emergence but insufficient to support vigorous subsequent elongation. In comparison, seedlings originating from laser-treated seeds under the same water condition displayed visibly greater root and shoot development and a more extensive root system. The seedling-growth measurements in Table 1 are consistent with these visual observations. At 1 ml, the mean shoot and root lengths of the laser-treated seedlings were 1.397 and 1.289 cm, respectively, compared with only 0.221 and 0.198 cm in the control group. Even at 0.5 ml, the treated seedlings maintained mean shoot and root lengths of 1.354 and 1.165 cm, whereas the corresponding control values decreased to 0.180 and 0.135 cm. The magnitude of these differences indicates that the effect of laser pretreatment extended beyond radicle emergence and improved early seedling establishment under restricted water availability. Because both groups received the same volume of water and were maintained under otherwise identical conditions, the enhanced growth of the treated seedlings suggests that laser pretreatment increased early seedling vigor and partially compensated for low water supply during the transition from germination to seedling establishment.

The improved germination and early growth of the laser-treated seeds may be associated with photobiological stimulation induced during the pretreatment stage. An appropriate optical dose may alter membrane properties, facilitate water uptake during subsequent imbibition, accelerate metabolic reactivation, and promote the mobilization of stored seed reserves.^20,64,65^ Blue-laser irradiation may also influence photoreceptor-mediated signaling and cellular redox regulation, potentially supporting the hormonal regulation, cell division, and cell elongation required for radicle and shoot development.^11,35,64^ Through these processes, laser-treated seeds may initiate germination more rapidly and use the limited available water more effectively during the transition from germination to seedling establishment.^20,66,67^ However, physiological, biochemical, and molecular parameters were not directly measured in the present study. These mechanisms should therefore be regarded as plausible explanations for the observed response rather than as experimentally confirmed pathways. Finally, Fig. 8 summarizes the overall response of lettuce seeds to progressive water limitation. As water availability decreased from 6 to 0.5 ml, the final germination percentage of the control seeds declined from approximately 72% to 57%. In contrast, the relative improvement produced by laser treatment increased from about 29% under standard water availability to nearly 46% under the most water-limited condition. This opposite trend indicates that the beneficial effect of laser irradiation became more pronounced as water availability decreased. Therefore, these findings suggest that optimized blue-laser treatment not only improves germination under standard laboratory conditions but also partially compensates for reduced water availability, supporting its potential as a sustainable physical seed-priming approach for improving early crop establishment under water-limited conditions.

**Figure 8:**
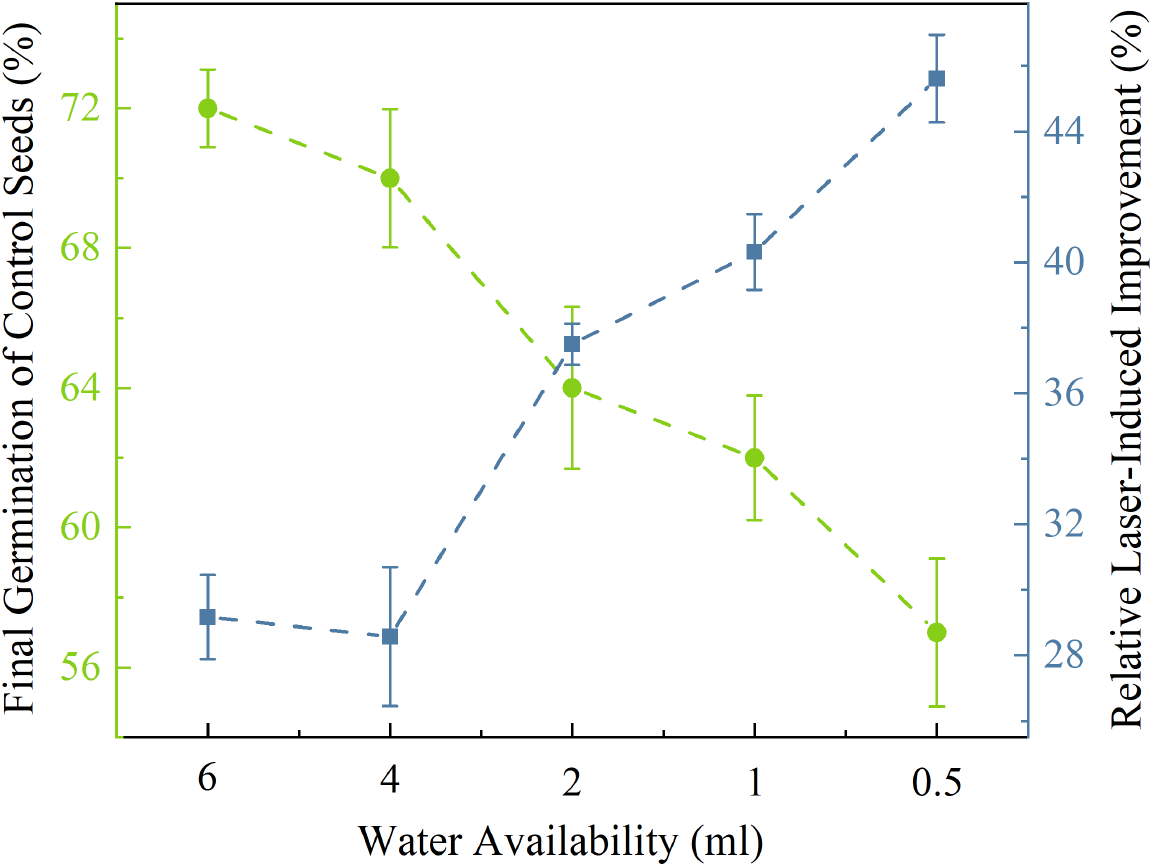
Effect of decreasing water availability on the final germination percentage of control lettuce seeds and the relative improvement induced by laser treatment. Green circles represent the final germination of control seeds, while blue squares indicate the relative laser-induced improvement compared with the corresponding control. Error bars represent the variability among three independent replicates.

## Conclusion

This study demonstrates that appropriately controlled irradiation with a 450 nm continuous-wave blue diode laser can enhance the germination and early seedling development of lettuce (*Lactuca sativa* L.). The response depended strongly on both laser power and exposure duration, supporting the existence of an optimal irradiation window. At a fixed exposure time of 2 min, increasing the laser power from 50 to 100 mW substantially improved germination performance, whereas further increases up to 250 mW produced only limited additional improvement, suggesting that the biostimulatory response approached saturation. Based on the overall germination and growth responses, 100 mW for 2 min was identified as the most effective treatment condition. Under this condition, the final germination percentage increased from approximately 65–70% in the non-irradiated control to approximately 90–95% in the laser-treated group. In addition to increasing final germination, the optimized laser treatment accelerated the main germination phase, increased the germination speed index, slightly improved the mean germination rate, and promoted early seedling development. Root elongation exhibited a stronger response than shoot elongation, indicating that the radicle and developing root tissues were particularly responsive to laser-induced biostimulation during early establishment. However, extending the irradiation duration beyond the optimum progressively reduced germination and seedling growth, indicating a transition from beneficial photobiostimulation to irradiation-induced stress. These findings emphasize that both laser power and exposure time must be optimized for each seed type, cultivar, and experimental condition.

Most importantly, laser pretreatment improved seed performance under reduced water availability. Across water volumes ranging from 0.5 to 6 ml per Petri dish, laser-treated seeds generally germinated earlier, maintained higher final germination percentages, and produced more developed seedlings than their corresponding controls. After 72 h, final germination ranged from approximately 83 to 93% in the laser-treated groups, compared with approximately 57 to 72% in the control groups. The relative improvement became more pronounced as water availability decreased, reaching its highest level under the most water-restricted condition. Thus, laser treatment did not eliminate the fundamental requirement for water during germination, but it partially compensated for low water supply by accelerating radicle emergence, shortening the time required to reach 50% germination, and supporting stronger early root and shoot establishment under restricted hydration. The comparison with sunlight further highlighted the advantage of controlled, wavelength-specific irradiation. Although equal-duration sunlight exposure produced a modest improvement over the control, blue laser treatment resulted in faster germination and a substantially higher final germination percentage. Moreover, experiments performed over 22 different days yielded a relative standard deviation of 8.65%, indicating a reasonably reproducible response despite the biological variability associated with seed age, storage conditions, and batch-to-batch differences. Therefore, these findings establish CW blue diode-laser irradiation as a promising contactless and chemical-free seed-priming strategy for enhancing lettuce germination, early seedling vigor, and establishment under reduced water availability. Because the water-limitation assay was performed in Petri dishes and physiological or molecular markers were not measured, future studies under greenhouse and field conditions should determine whether these early benefits persist during later developmental stages and should evaluate treatment scalability, energy efficiency, and applicability to other cultivars and crop species.

## Acknowledgement

This work was supported by TUBITAK projects nos. 124F388 and 126F048.

## TOC Graphic

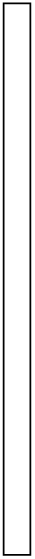

